# Developmental programming: Transcriptional regulation of visceral and subcutaneous adipose by prenatal bisphenol-A in female sheep

**DOI:** 10.1101/2019.12.12.874446

**Authors:** John F. Dou, Muraly Puttabyatappa, Vasantha Padmanabhan, Kelly M. Bakulski

**Affiliations:** Department of Epidemiology, School of Public Health, University of Michigan, Ann Arbor, Michigan, USA; Department of Pediatrics, University of Michigan, Ann Arbor, Michigan, USA

**Keywords:** Adipose depots, Bisphenol A, Endocrine disruptor, RNA sequencing

## Abstract

**Background:** Bisphenol-A (BPA) exposure is widespread and early life exposure is associated with metabolic syndrome. While visceral adipose tissue (VAT) and subcutaneous adipose tissue (SAT) are implicated in the development of metabolic syndrome, the adipose depot-specific effects of prenatal BPA treatment are poorly understood.

**Objective:** To determine the impact of prenatal BPA exposure on the transcriptome of VAT and SAT adipose depots.

**Methods:** RNA sequencing was performed on SAT and VAT from 21-month old control and prenatal BPA-treated female sheep. Differences in transcriptional profiling of SAT and VAT in controls and the effect of prenatal BPA treatment on individual genes and gene pathways were determined.

**Results:** There were 179 differentially expressed genes (p_adjusted_<0.05, log_2_-fold change >2.5) between SAT and VAT. Development and immune response pathways were upregulated in SAT, while metabolic pathways were upregulated in VAT. In SAT, BPA-treatment resulted in differential expression of 108 genes (78% upregulated with BPA) and altered pathways (immune response downregulated, RNA processing upregulated). In contrast in VAT, BPA-treatment differentially expressed 4 genes and upregulated chromatin and RNA processing pathways.

**Conclusion:** Prenatal BPA-treatment induces adult depot-specific alterations in RNA expression in inflammation, RNA processing, and chromatin, reflecting the diverse roles of SAT and VAT in regulating lipid storage and insulin sensitivity. These adipose tissue transcriptional dysregulations may contribute to the metabolic disorders observed in prenatal BPA-treated female sheep.

## Introduction

The worldwide prevalence of obesity tripled between 1975 and 2016, until 650 million adults were classified as obese and another 1.25 billion adults were overweight (WHO, 2018). This rapid increase in obesity and associated complications created an annual economic burden due to healthcare costs estimated between $147 and $210 billion in the United States (US) alone (Levi et al., 2010; Regnier and Sargis, 2014). The main risk factors for the development of obesity are genetic predisposition and lifestyle factors, including increased caloric intake and decreased energy expenditure (Williams et al., 2015). However, these factors alone fail to explain the global rise in obesity prevalence. Accumulating evidence from epidemiological and animal studies demonstrate early development exposures to environmental endocrine disrupting chemicals, such as bisphenol A (BPA), are risk factors for obesity (Zimmet and Alberti, 2006; Holtcamp, 2012; Kelishadi et al., 2013; Heindel et al., 2015). Over 90% of US adults and 96% of pregnant women have detectable urinary concentrations of BPA (Calafat et al., 2008; Woodruff et al., 2011), indicating widespread exposure and risk particularly during the prenatal period. Used in the production of epoxy and polycarbonate resins, BPA is found in common household plastics. Although its main endocrine disrupting function is its estrogenic activity (Vogel, 2009; Vom Saal, 2016), BPA is also known for its obesogenic role (Legeay and Faure, 2017). BPA affects adipose tissue via estrogenic actions as well as disruption of the thyroid receptor and activation of peroxisome proliferator-activated receptor gamma (PPARG) (Vom Saal et al., 2012; Ahmed, 2016; Ahmed and Atlas, 2016; Vom Saal, 2016). Because obesity is a global public health challenge and early life exposure to BPA may be obesogenic there is a need to understand the underlying mechanisms of BPA actions.

Many human epidemiological studies have linked BPA exposure to metabolic defects, such as insulin resistance, diabetes, obesity and metabolic syndrome (Newbold, 2010; Teppala et al., 2012; Wang et al., 2012). There is suggestive evidence linking developmental BPA exposure with reduced birth weight (Veiga-Lopez et al., 2015a), a major risk factor for adult onset cardio-metabolic disorders (Barker, 2005). Animal studies provide direct evidence of developmental BPA exposure induced metabolic defects (vom Saal et al., 2007; Alonso-Magdalena et al., 2010; Veiga-Lopez et al., 2016; Wassenaar et al., 2017), where early-life exposure to BPA is significantly and positively associated with obesity-related adipose tissue disruptions (Wassenaar et al., 2017). In the sheep, the animal model used in this study, prenatal BPA-treated female offspring develop metabolic defects, manifested as peripheral insulin resistance with compensatory hyperinsulinemia and adipose tissue disruptions (Veiga-Lopez et al., 2015b; Veiga-Lopez et al., 2016). These sheep develop adipocyte hypertrophy and visceral adipose tissue (VAT) and subcutaneous adipose tissue (SAT) specific changes in markers of inflammation, oxidative stress and adipose differentiation markers (Veiga-Lopez et al., 2015b; Veiga-Lopez et al., 2016). Although both VAT and SAT are predominantly white adipose depots, they are morphologically and functionally different. The VAT depot is more responsive to the catecholamine-induced lipolysis and less responsive to the antilipolytic effect of insulin and therefore have a higher rate of fatty acid turnover and lipolysis (Engfeldt and Arner, 1988; Ibrahim, 2010). In contrast, SAT adipocytes are responsive to insulin action and favor uptake and storage of free fatty acids and triglycerides. In addition, these depots also exhibit a distinct pattern of gene expression such as increased expression of proinflammatory cytokines and adipokines such as tumor necrosis factor alpha (TNF), C reactive peptide (CRP) and interleukin 6 (IL6) in VAT than SAT (Ibrahim, 2010). Because of these functional and transcriptional differences, accumulation of VAT is a risk factor for development of cardiometabolic defects (Lemieux and Despres, 1994; Bjorntorp, 2000; Dobbelsteyn et al., 2001). Therefore, understanding the depot-specific effects of prenatal BPA treatment can aid in delineating the mechanism of development of adipose and metabolic defects, which are poorly understood.

In the adipose tissues, the effects of BPA are mediated at least in part through activation of nuclear transcription factors – PPARG and steroidal hormone receptors (Wetherill et al., 2007; Boucher et al., 2014a; Ahmed, 2016; Ahmed and Atlas, 2016). Activation of these receptors are known to influence various aspects of adipose tissue function including adipogenesis, adipocyte proliferation and differentiation, and glucose and lipid metabolism (Moreno-Navarrete and Fernández-Real, 2017; Newell-Fugate, 2017; Corrales et al., 2018) via changes in patterns of gene expression (Polvani et al., 2016; Mota de Sa et al., 2017). Various gene families influenced by BPA exposure in adipose tissue include adipokines – adiponectin, leptin, inflammatory cytokines – IL6 and TNF, mediators of lipogenesis - Lipoprotein lipase (LPL) and Glycerol-3-phosphate dehydrogenase (GPDH) and adipogenesis – PPARG, LPL and fatty acid-binding protein 4 (FABP4) (Masuno et al., 2005; Ben-Jonathan et al., 2009; Melzer et al., 2011). In sheep, prenatal BPA treatment altered the expression of adiponectin, oxidative stress markers, steroidogenic enzymes and steroid receptors in a dose-dependent manner in the VAT (Puttabyatappa et al., 2019). Treatment also induced depot specific changes in the expression of adiponectin and macrophage marker in VAT and SAT (Veiga-Lopez et al., 2016). Considering that VAT and SAT have different roles in the maintenance of metabolic homeostasis (Zhang et al., 2014) and BPA influences transcription factor regulation, understanding the depot-specific gene expression disruptions induced by BPA exposure are of interest.

As described above, BPA affects adipose tissue function through divergent means, thus creating a challenge to gain a comprehensive understanding of the signaling pathways underlying these functional outcomes. High throughput quantitative transcriptome profiling using whole adipose tissue RNA-sequencing (RNAseq) is used to identify depot-specific gene and gene networks disrupted by prenatal BPA exposure. The major advantage of this technique is examining the whole cell/tissue transcriptome profile and unlike with microarray technology, prior probe selection does not depend on genome annotation, avoiding the related biases introduced during hybridization of microarrays (Wang et al., 2009; Zhao et al., 2014). In addition, it also provides the ability through bioinformatic analysis, identification of gene networks affected (Wang et al., 2009). Therefore, the goals of the present study were 1) to compare transcriptional differences between VAT and SAT depots, 2) determine genes and gene networks disrupted by prenatal BPA treatment and 3) to identify potential pathways that underlie lasting adipose-depot specific functional changes stemming from developmental exposure to BPA.

## METHODS

### Animals

All animal procedures were approved by the Institutional Animal Care and Use Committee of the University of Michigan and are consistent with the National Institutes of Health’s Guide for the Care and Use of Laboratory Animals. These studies were conducted at the University of Michigan Sheep Research Facility (Ann Arbor, MI) using the Suffolk sheep breed acquired from local farmers. The maintenance, breeding, prenatal treatments and lambing were performed as described previously (Manikkam et al., 2004). All animals including the control group were maintained together and any exposures to potential sources of phytoestrogens via diet were therefore similar across control and prenatal BPA treated groups.

### Prenatal BPA Treatment

Pregnant sheep were randomly assigned to control (C) and BPA groups. Control mothers received only the vehicle (corn oil) and BPA animals received 0.5 mg/kg /day of BPA (purity ≥99%, cat. no. 239658; Aldrich Chemical, Milwaukee, WI) dissolved in corn oil. Both treatments were administered daily through subcutaneous injections from days 30 through 90 of gestation (term: ∼147 days). This BPA dosing resulted in 2.62±0.52ng/ml of free BPA umbilical artery on day 90 of fetal life (Veiga-Lopez et al., 2013a), which are within the range found in mid-gestation umbilical cord blood concentrations of free BPA (Gerona et al., 2013; Veiga-Lopez et al., 2015b; Lee et al., 2018). The female lambs were weaned at ∼8 weeks of age and provided with a maintenance diet consisting of 0.64 kg of corn, 0.64 kg hay·lamb-1·day-1, and 0.014 kg of supplement (36% crude protein) to prevent the development of obesity. Wooden feeders with crossover metallic bars were used. Only one female offspring from each dam was utilized if twin pregnancies were involved. The effects of prenatal BPA exposure on ovarian follicular dynamics, LH surge, insulin sensitivity, adiposity and insulin sensitivity mediators utilizing the animals from this cohort have been previously reported (Veiga-Lopez et al., 2013b; Veiga-Lopez et al., 2014; Veiga-Lopez et al., 2015b; Puttabyatappa et al., 2019).

### Tissue Collection

Both VAT and SAT were collected during the second breeding season at ∼21 months of age. Before collection, estrus was synchronized with two injections of prostaglandin F2α (PGF2α, 10 mg, i.m.; Lutalyse, Pfizer Animal Health, Florham Park, NJ) administered 11 days apart. Tissues were harvested 24 hours after the second PGF2α injection during the follicular phase to maintain comparable steroid environment. All animals were euthanized by barbiturate overdose (Fatal Plus; Vortech Pharmaceuticals, Dearborn, MI) and VAT from the mesenteric fat surrounding the ventral sac of the rumen and SAT from the sternal region were obtained. Tissues were flash-frozen and stored at -80 °C until processed.

### RNA Sequencing

The University of Michigan Advanced Genomics and Next Generation Sequencing Core prepared cDNA libraries and performed RNA sequencing on VAT and SAT tissues from randomly selected control and prenatal BPA-treated animals (n=4 / treatment group). Libraries were prepared with SMARTer universal low input RNA kit (Takara, Mountain View, CA) following ribosome depletion. Libraries were split across four lanes and sequenced on the HiSeq NovaSeq 6000 platform (Illumina, San Diego, CA). Single end sequencing was performed for 76 cycles. Raw data from this experiment are available at the Genome Expression Omnibus (GEO, accession number pending).

### Data Processing and Quality Control

Raw fastq files were examined using fastQC (version 0.11.5) (Andrews, 2010). Reports generated for the 16 samples across 4 lanes were summarized using multiQC (version 0.9) (Ewels et al., 2016). Across samples and at all base positions, mean quality scores were high. GC content ranged from 49% to 58%. There were between 32% and 64% of reads duplicated. Reads were mapped to a sheep reference genome (Oar_rambouillet_v1.0) using the Spliced Transcripts Alignment to a Reference (STAR) (version 2.6.0c) program (Dobin et al., 2013). Post alignment, QoRTs (version 1.3.6) (Hartley and Mullikin, 2015) was used to examine further quality control metrics. Sample distributions in quality metrics were clustered by the subcutaneous and visceral adipose tissue types. Alignment soft clipping rate was highest at the start and end of read cycles, with a larger spike of approximately 30-40% at the start. Approximately 15-20% of reads per sample were dropped for multi-mapping. A large proportion of reads (approximately 50-70%) mapped to non-gene regions, which may reflect lack of annotation in the sheep genome. Following mapping, featureCounts (version 1.6.1) (Liao et al., 2014) was used to quantify aligned reads. Default behavior was used to drop multi-mapping reads and count features mapping to exons.

### Differential Expression Analysis

First, aligned and quantified reads were analyzed in R statistical software (version 3.6.0) with the DESeq2 package (version 1.24.0) (Love et al., 2014). Lanes were collapsed to single samples. Default settings for DESeq2 were used for filtering of genes with low normalized mean counts. Principal components analysis was performed, calculated on variance stabilizing transformed values of the expression data. Principal components were plotted to examine clustering by sample type.

Differentially expressed genes were examined by tissue type (VAT versus SAT) and by treatment (control versus BPA). To investigate differences by tissue type, samples from SAT and VAT of control animals were compared. To investigate the effect of BPA, the BPA-treated group was compared to the control, separately for SAT and VAT. Volcano plots of results were created using the EnhancedVolcano (version 1.2.0) package, after applying log fold change shrinkage using the “apeglm” prior (Zhu et al., 2019). To account for multiple comparisons, an adjusted p-value < 0.05 was set as threshold. To prioritize genes for further investigation, an absolute log_2_ fold-change > 2.5 in the tissue type comparison, and an absolute log_2_ fold-change > 1.5 in BPA treatment comparisons were considered.

### Gene Set Enrichment

Enriched gene set terms were tested using RNA-enrich (Lee et al., 2016). Sheep genes were annotated to human orthologs using Ensembl identifiers and the BioMart package (Durinck et al., 2009). Three pathway analyses were performed using the directional RNA-Enrich test for 1) the tissue type comparison, 2) for BPA exposure in SAT, and 3) for BPA exposure in VAT. RNA-Enrich does not require a significance cutoff for genes, and accounts for the relationship between gene read count and statistical significance. We ran tests with the Biocarta Pathway, EHMN metabolic pathways, Gene Ontology, KEGG Pathway, Panther Pathway, and transcription factors databases selected. Maximum concept size was set to 1000 genes, we left other options at default values.

### Network Analysis

Gene interaction networks were visualized using the STRING database of protein-protein interactions (Franceschini et al., 2013). Genes highlighted were contained in one of two pathways: inflammatory response and chromatin modification. Genes with p<0.05 in one or both fat deposits in relation to BPA exposure were included in network diagrams. A score threshold of 300 was set for inclusion of gene interactions.

Code to conduct the analyses presented in this manuscript are available through GitHub (https://github.com/bakulskilab).

## Results

### Descriptive Statistics

The study consisted of two adipose tissue types and two treatment groups. Each adipose depot and treatment combination had four samples. Following alignment and quantification, samples had between 3,046,126 to 14,305,764 reads assigned to features numbering from 16,926 to 18,914 (**Supplemental Table 1**). In principal component plots, clustering by adipose tissue type and treatment group were observed (**Supplemental Figure 1**).

### Transcriptional Differences Between VAT and SAT Depots

Following filtering of genes with low normalized mean counts and extreme count outliers, 17,242 genes were analyzed. There were 2,196 genes differentially expressed (adjusted p<0.05), 180 genes with a log_2_-fold change magnitude difference > 2.5, and 179 genes met both criteria (**Figure 1A**). Of the genes meeting these statistical and magnitude criteria, 61% (109 genes) had higher expression in SAT. Top differentially expressed genes in SAT relative to VAT by adjusted p-value include epoxide hydrolase 2 (*EPHX2*) (−2.61 lower log_2_-fold change expression, adjusted-p=9.1×10^−37^), homeobox C9 (*HOXC9*) (3.7 higher log_2_-fold change expression, adjusted-p=2.9×10^−31^) (**Supplemental Figure 2A**), transferrin (*TF*) (−2.8 lower log_2_-fold change expression, adjusted-p=1.4×10^−27^) and cut like homeobox 1 (*CUX1*) (−1.2 lower log_2_-fold change expression, adjusted-p=8.9×10^−27^) (**Table 1, Supplemental Table 2**). Top genes ordered by shrunk log_2_-fold change values were Iroquois homeobox 2 (*IRX2*) (12.1 higher log_2_-fold change, p-adjusted=1.7×10^−19^) (**Supplemental Figure 2B**), leucine rich repeat neuronal 4 (*LRRN4*) (−10.9 lower log_2_-fold change, adjusted-p=1.3×10^−8^), *HOXA9* (10.0 higher log_2_-fold change, adjusted-p=1.2×10^−13^) and transcription factor 21 (*TCF21*) (−9.7 lower log_2_-fold change, adjusted-p=3.2×10^−12^). Compared to VAT, SAT showed upregulation in genes that participate in skeletal system morphogenesis pathways, and downregulation of genes involved in mitochondrial membrane and vasodilation pathways (**Supplemental Table 3**). A total of 260 pathways had FDR<0.05, with a set of pathways upregulated in SAT were involved in cellular and tissue development, and immune response, while pathways involved in metabolism, mitochondria, homeostasis, transcription, and translation were downregulated in SAT (**Figure 1B**).

**Table 1.** Differential gene expression between subcutaneous adipose tissue (SAT) and visceral adipose tissue (VAT) depots from 21-month old female sheep.

| Gene | Base Mean Expression | Log<sub>2</sub>(Fold change in expression SAT versus VAT) | Standard error | p-value | p-adjusted |
| --- | --- | --- | --- | --- | --- |
| EPHX2 | 711.9 | -2.61 | 0.198 | 5.30E-41 | 9.13E-37 |
| HOXC9 | 72.3 | 3.73 | 0.307 | 3.39E-35 | 2.92E-31 |
| TF | 911.03 | -2.84 | 0.25 | 2.44E-31 | 1.40E-27 |
| CUX1 | 1376.11 | -1.15 | 0.103 | 2.07E-30 | 8.92E-27 |
| BARX1 | 97.64 | -5.18 | 0.461 | 4.31E-30 | 1.48E-26 |
| ZFHX4 | 1888.61 | 1.14 | 0.104 | 3.88E-29 | 1.12E-25 |
| PITX2 | 100.64 | 8.36 | 0.785 | 3.87E-26 | 9.54E-23 |
| LOC105614720 | 37.93 | 3.59 | 0.352 | 1.87E-25 | 3.75E-22 |
| SLIT2 | 791.71 | -3.7 | 0.364 | 1.96E-25 | 3.75E-22 |
| LOC106991591 | 51.69 | 2.72 | 0.269 | 3.28E-25 | 5.66E-22 |
| IRX2 | 99.5 | 12.1 | 2.85 | 1.08E-22 | 1.69E-19 |
| EN1 | 127.24 | 4.37 | 0.476 | 2.04E-21 | 2.94E-18 |
| TMEM120A | 282.55 | -1.69 | 0.19 | 3.79E-20 | 5.03E-17 |
| MPPED2 | 42.57 | 3.65 | 0.411 | 4.92E-20 | 6.06E-17 |
| RBPMS2 | 152.81 | -1.8 | 0.208 | 3.26E-19 | 3.74E-16 |
| GPAT3 | 276.56 | -3.54 | 0.415 | 8.12E-19 | 8.75E-16 |
| GPC3 | 1534.41 | 1.73 | 0.209 | 1.06E-17 | 1.08E-14 |
| HOXA9 | 117.69 | 10 | 1.45 | 1.22E-16 | 1.17E-13 |
| TBX15 | 135.54 | 3.13 | 0.397 | 1.49E-16 | 1.35E-13 |

**Figure 1.**
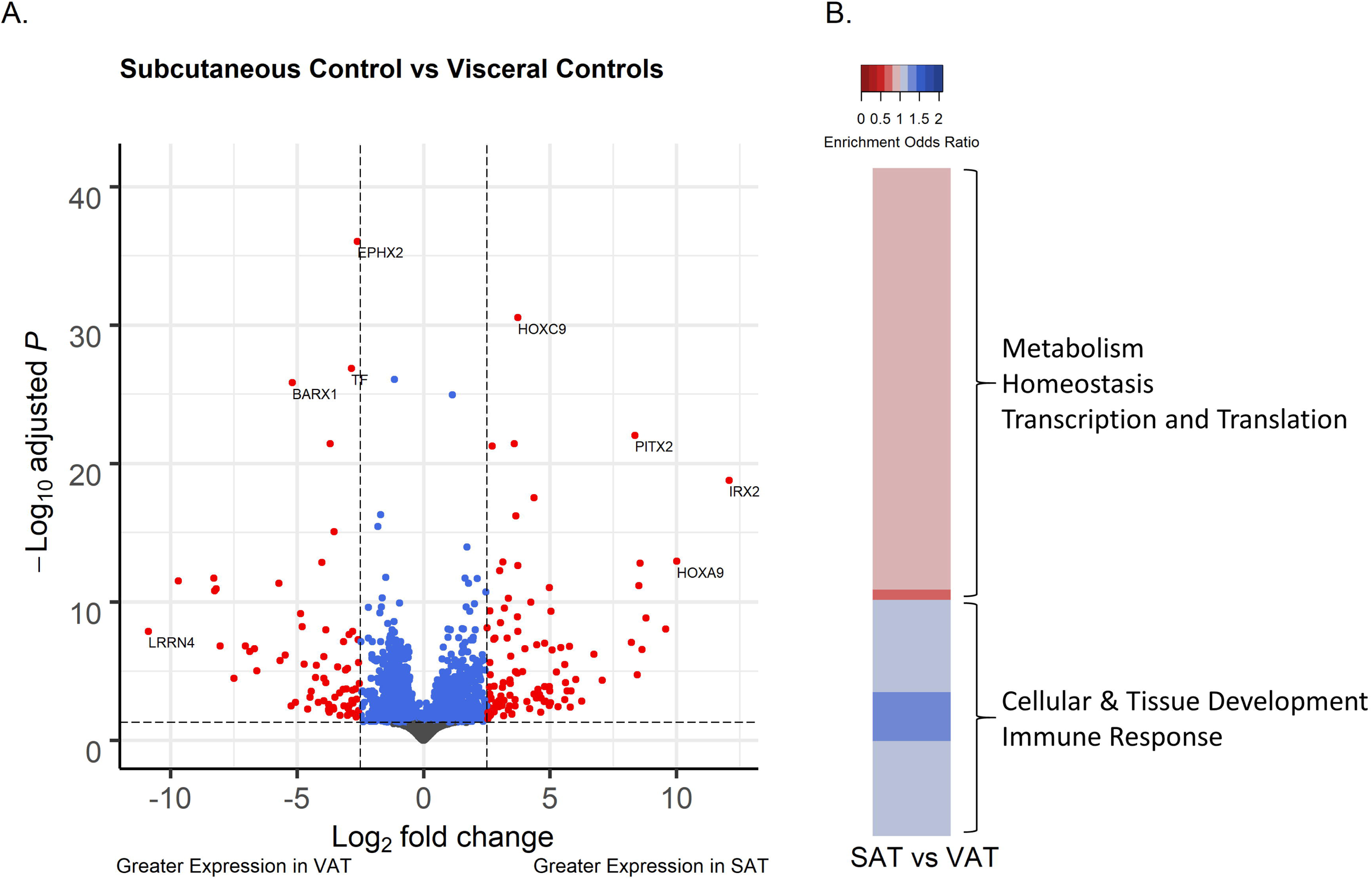
Comparison of the gene expression of 21-month old female sheep in subcutaneous adipose tissue (SAT) relative to visceral adipose tissue (VAT). (**A**) Volcano plot of differential gene expression by adipose tissue type, comparing the SAT versus VAT from control sheep. Genes are plotted by log-fold change, using “apeglm” for shrinkage, and -log adjusted p-values. Blue points have adjusted p-values < 0.05. Red points also have log-fold change > 2.5. (**B**) Heatmap of pathways with FDR < 0.05 in the comparison of SAT versus VAT tissue differences. Upregulated gene sets are in blue, downregulated gene sets in red.

### Transcriptional Differences Between Control and BPA-Treated Adipose Depots

In SAT, after filtering for low normalized mean counts, 16,388 genes remained in analysis. There were 319 genes differentially expressed between control and BPA-treated groups at a false discovery rate adjusted p-value < 0.05, 118 genes differentially expressed with an absolute log_2_-fold change magnitude of difference > 1.5, and 108 genes met both these conditions (**Figure 2A**). Of the genes meeting these statistical and magnitude criteria, 78% (84 genes) had higher expression in BPA-treated group. Ordered by adjusted p-value, the top differentially expressed genes with BPA treatment relative to control treatment include uncharacterized gene (*LOC114116052*) (−0.006 lower log_2_-fold change expression adjusted-p=7.7×10^−9^), cyclin N-terminal domain containing 2 (*CNTD2*) (1.9 higher log_2_-fold change expression, adjusted-p=7.7×10^−9^) (**Supplemental Figure 3A**), 5S ribosomal RNA *LOC114116728* (2.5 higher log_2_-fold change expression, adjusted-p=5.9×10^−8^), and *LOC114116239* (2.8 higher log_2_-fold change expression, adjusted-p=2.3×10^−7^) (**Table 2, Supplemental Table 4**). Ordered by log_2_-fold change, top genes include U6 spliceosomal RNA *LOC114117140* (3.9 higher log_2_-fold change expression, adjusted-p=2.2×10^−4^), 5S ribosomal RNA *LOC114112289* (3.6 higher log_2_-fold change expression, adjusted-p=2.3×10^−7^) (**Supplemental Figure 3B**), small nucleolar RNA SNORD61 *LOC114111819* (3.5 higher log_2_-fold change expression, adjusted-p=0.01), and troponin C2 (*TNNC2*) (3.4 higher log_2_-fold change expression, adjusted-p=0.02). Hierarchical clustering based on gene expression profiles show clear clustering in SAT samples by BPA treatment (**Supplementary Figure 4**).

**Table 2.**
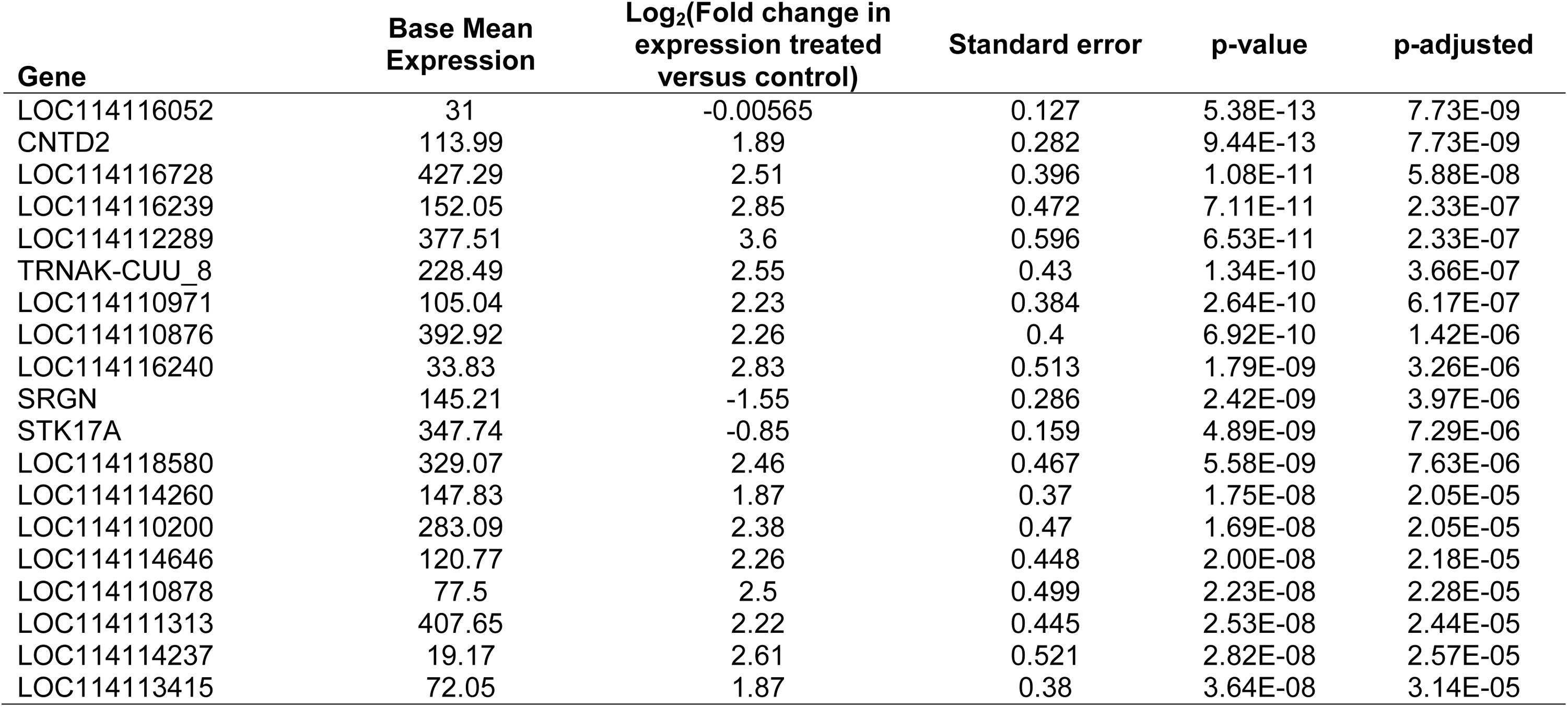
Prenatal BPA treatment induced differential gene expression in the subcutaneous adipose tissue depot in 21-month old female sheep.

| Gene | Base Mean Expression | Log<sub>2</sub>(Fold change in expression treated versus control) | Standard error | p-value | p-adjusted |
| --- | --- | --- | --- | --- | --- |
| LOC114116052 | 31 | -0.00565 | 0.127 | 5.38E-13 | 7.73E-09 |
| CNTD2 | 113.99 | 1.89 | 0.282 | 9.44E-13 | 7.73E-09 |
| LOC114116728 | 427.29 | 2.51 | 0.396 | 1.08E-11 | 5.88E-08 |
| LOC114116239 | 152.05 | 2.85 | 0.472 | 7.11E-11 | 2.33E-07 |
| LOC114112289 | 377.51 | 3.6 | 0.596 | 6.53E-11 | 2.33E-07 |
| TRNAK-CUU_8 | 228.49 | 2.55 | 0.43 | 1.34E-10 | 3.66E-07 |
| LOC114110971 | 105.04 | 2.23 | 0.384 | 2.64E-10 | 6.17E-07 |
| LOC114110876 | 392.92 | 2.26 | 0.4 | 6.92E-10 | 1.42E-06 |
| LOC114116240 | 33.83 | 2.83 | 0.513 | 1.79E-09 | 3.26E-06 |
| SRGN | 145.21 | -1.55 | 0.286 | 2.42E-09 | 3.97E-06 |
| STK17A | 347.74 | -0.85 | 0.159 | 4.89E-09 | 7.29E-06 |
| LOC114118580 | 329.07 | 2.46 | 0.467 | 5.58E-09 | 7.63E-06 |
| LOC114114260 | 147.83 | 1.87 | 0.37 | 1.75E-08 | 2.05E-05 |
| LOC114110200 | 283.09 | 2.38 | 0.47 | 1.69E-08 | 2.05E-05 |
| LOC114114646 | 120.77 | 2.26 | 0.448 | 2.00E-08 | 2.18E-05 |
| LOC114110878 | 77.5 | 2.5 | 0.499 | 2.23E-08 | 2.28E-05 |
| LOC114111313 | 407.65 | 2.22 | 0.445 | 2.53E-08 | 2.44E-05 |
| LOC114114237 | 19.17 | 2.61 | 0.521 | 2.82E-08 | 2.57E-05 |
| LOC114113415 | 72.05 | 1.87 | 0.38 | 3.64E-08 | 3.14E-05 |

**Figure 2.**
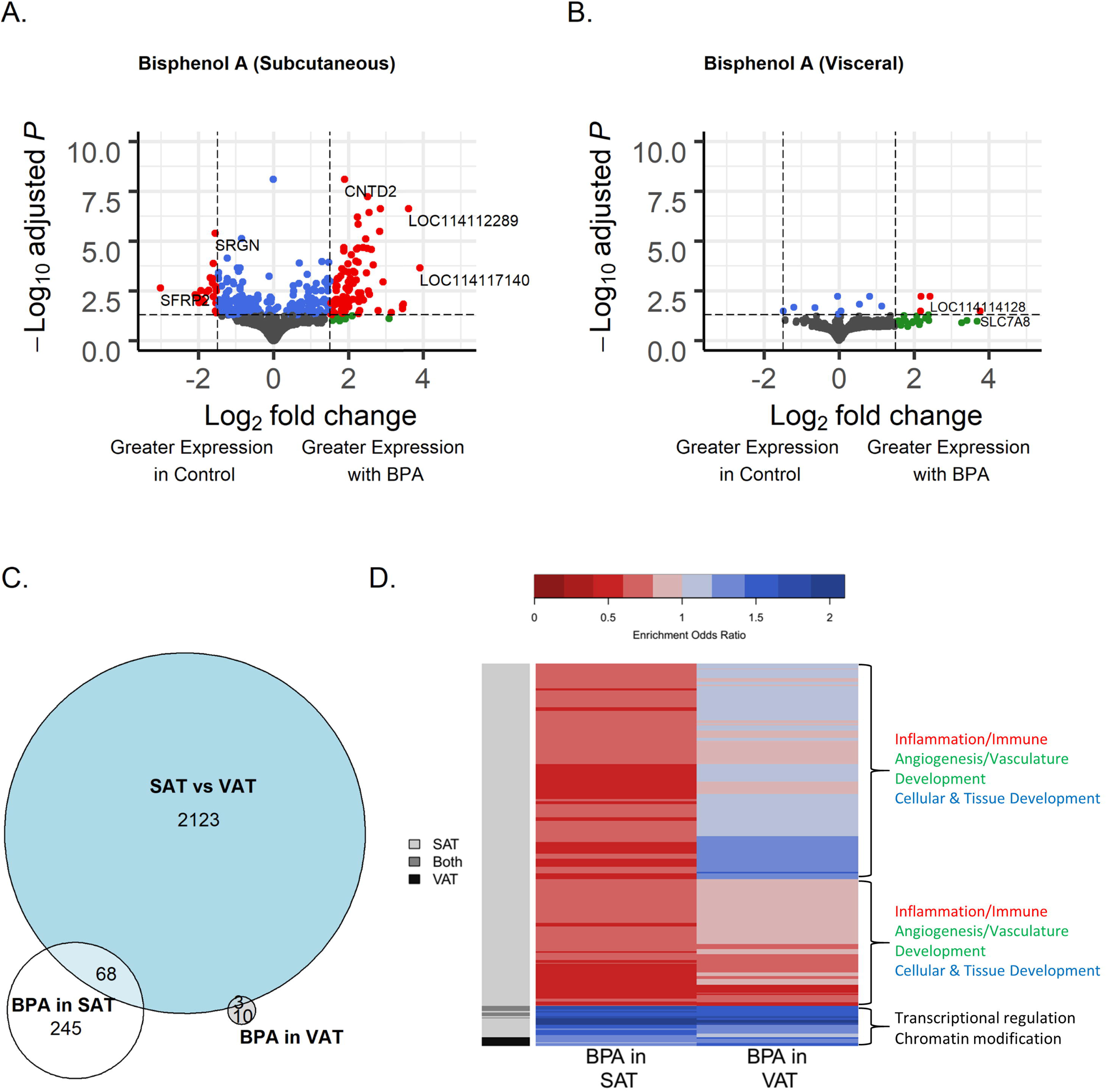
Comparison of the gene expression of 21-month old female sheep in subcutaneous adipose tissue from prenatal bisphenol-A (BPA) treated and control t groups. (**A**) Volcano plot of differential gene expression by prenatal BPA treatment in subcutaneous adipose tissue (SAT). (**B**). Volcano plot of differential gene expression by prenatal BPA treatment in visceral adipose tissue (VAT). Genes are plotted by log-fold change, using “apeglm” for shrinkage, and -log adjusted p-values. Blue points have adjusted p-values < 0.05. Green points have shrunk log fold change > 1.5. Red points have both adjusted p-value < 0.05 and log fold change > 1.5. (**C**). Venn diagram of number of FDR significant genes in tissue comparison, BPA effect in SAT, and BPA effect in VAT. (**D**). Heatmap of clustering of significant genes sets from RNA-Enrich with FDR < 10^−6^ in either the BPA comparison in SAT or the BPA comparison in VAT. Upregulated gene sets are in blue, downregulated gene sets in red.

In VAT, following filtering of genes with low normalized mean counts and extreme count outliers, 15,537 genes were analyzed. Comparing BPA treated and control VAT, there were 13 genes differentially expressed at a false discovery rate adjusted p-value<0.05, 27 genes differentially expressed with an absolute log_2_-fold change magnitude of difference > 1.5, and 4 genes met both criteria (**Figure 2B**). These 4 genes are small nucleolar RNA SNORA61 *LOC114113403* (2.2 higher log_2_-fold-change expression with BPA treatment, adjusted-p=0.006), uncharacterized gene (*LOC114114128)* (2.4 higher log_2_-fold-change expression, adjusted-p=0.006), solute carrier family 7 member 8 (*SLC7A8*) (3.8 higher log_2_-fold change expression, adjusted-p=0.03), and *LOC114110120* (2.2 higher log_2_-fold change expression, adjusted-p=0.03) (**Table 3, Supplemental Table 5**). The number of genes impacted by BPA in VAT was smaller than the effects in SAT, and there was no overlap in FDR significant genes between the two tissue types (**Figure 2C**).

**Table 3.** Prenatal BPA treatment induced differential gene expression in the visceral adipose tissue depot in 21-month old female sheep.

| Gene | Base Mean Expression | Log<sub>2</sub>(Fold change in expression treated versus control) | Standard error | p-value | p-adjusted |
| --- | --- | --- | --- | --- | --- |
| LOC114113403 | 198.28 | 2.18 | 0.518 | 9.24E-07 | 0.00598 |
| LOC114114128 | 33.97 | 2.43 | 0.55 | 3.95E-07 | 0.00598 |
| ARID3A | 371.95 | 0.813 | 0.193 | 1.16E-06 | 0.00598 |
| LOC114116716 | 9.1 | -0.0325 | 0.112 | 1.59E-06 | 0.00618 |
| HDGF | 859.79 | 0.543 | 0.139 | 4.85E-06 | 0.0151 |
| FUK | 195.13 | 1.14 | 0.305 | 7.30E-06 | 0.0189 |
| PSTPIP2 | 169.07 | -1.2 | 0.328 | 9.82E-06 | 0.0218 |
| DDR2 | 1958.64 | -0.645 | 0.176 | 1.16E-05 | 0.0225 |
| SLC7A8 | 21.55 | 3.76 | 1.12 | 2.26E-05 | 0.0329 |
| HTR1B | 23.37 | -1.49 | 0.435 | 2.54E-05 | 0.0329 |
| ANKRD55 | 12.08 | 0.0538 | 0.126 | 2.24E-05 | 0.0329 |
| LOC114110120 | 33.89 | 2.18 | 0.645 | 2.45E-05 | 0.0329 |
| LOC105605961 | 25.55 | -0.0213 | 0.107 | 3.91E-05 | 0.0468 |
| LOC114117716 | 290.32 | 2.39 | 0.75 | 4.66E-05 | 0.0517 |
| LOC105616380 | 144.55 | 1.37 | 0.435 | 5.51E-05 | 0.0571 |
| LOC114116238 | 212.91 | 1.73 | 0.554 | 5.92E-05 | 0.0575 |
| RECQL4 | 15.58 | 2.07 | 0.671 | 6.76E-05 | 0.0601 |
| ZNF565 | 85.13 | -0.605 | 0.191 | 7.21E-05 | 0.0601 |
| CARMIL1 | 397.35 | 0.532 | 0.167 | 7.35E-05 | 0.0601 |

Several up and downregulated pathways in relation to differential gene expression by BPA were identified through RNA-Enrich. In SAT, BPA downregulated several immune cell pathways (e.g. leukocyte activation, inflammatory response, cytokine production) and upregulated RNA processing pathways (RNA splicing, mRNA metabolic process) (**Figure 2D**; **Supplemental Table 6**). BPA in VAT was associated with upregulation of chromatin modification pathway and RNA processing pathways, such as mRNA processing (**Figure 2D; Supplemental Table 7**). Transcriptional regulation and chromatin modification pathways were upregulated in response to BPA in both SAT and VAT (**Figure 2D**). Inflammation, immune, angiogenesis/vasculature development, and cellular and tissue development related pathways were downregulated in SAT, with more mixed or smaller impacts in VAT (**Figure 2D)**.

The inflammatory response pathway had more genes affected by BPA in SAT than in VAT (**Figure 3**). Several well-connected genes with many interactions were impacted by BPA in SAT (toll like receptor 2 [*TLR2*], MYD88 innate immune signal transduction adaptor [*MYD88*], mitogen-activated protein kinase 14 [*MAPK14*]). The chromatin modification pathway had many genes in SAT and VAT impacted by BPA, with several overlapping genes affected by BPA in both cell types (**Figure 4**). Genes with high degree of interactions had expression modified by BPA in VAT, including histone deacetylase 1 (*HDAC1*), DNA methyltransferase 1 (*DNMT1*), cAMP responsive element binding protein binding protein (*CREBBP*), euchromatic histone lysine methyltransferase 1 (*EHMT1*), and *EHMT2*. There were also highly connected genes with BPA effects in both SAT and VAT (DNA methyltransferase 1 associated protein 1 [*DMAP1*], SWI/SNF related, matrix associated, actin dependent regulator of chromatin, subfamily a, member 4 [*SMARCA4*]).

**Figure 3.**
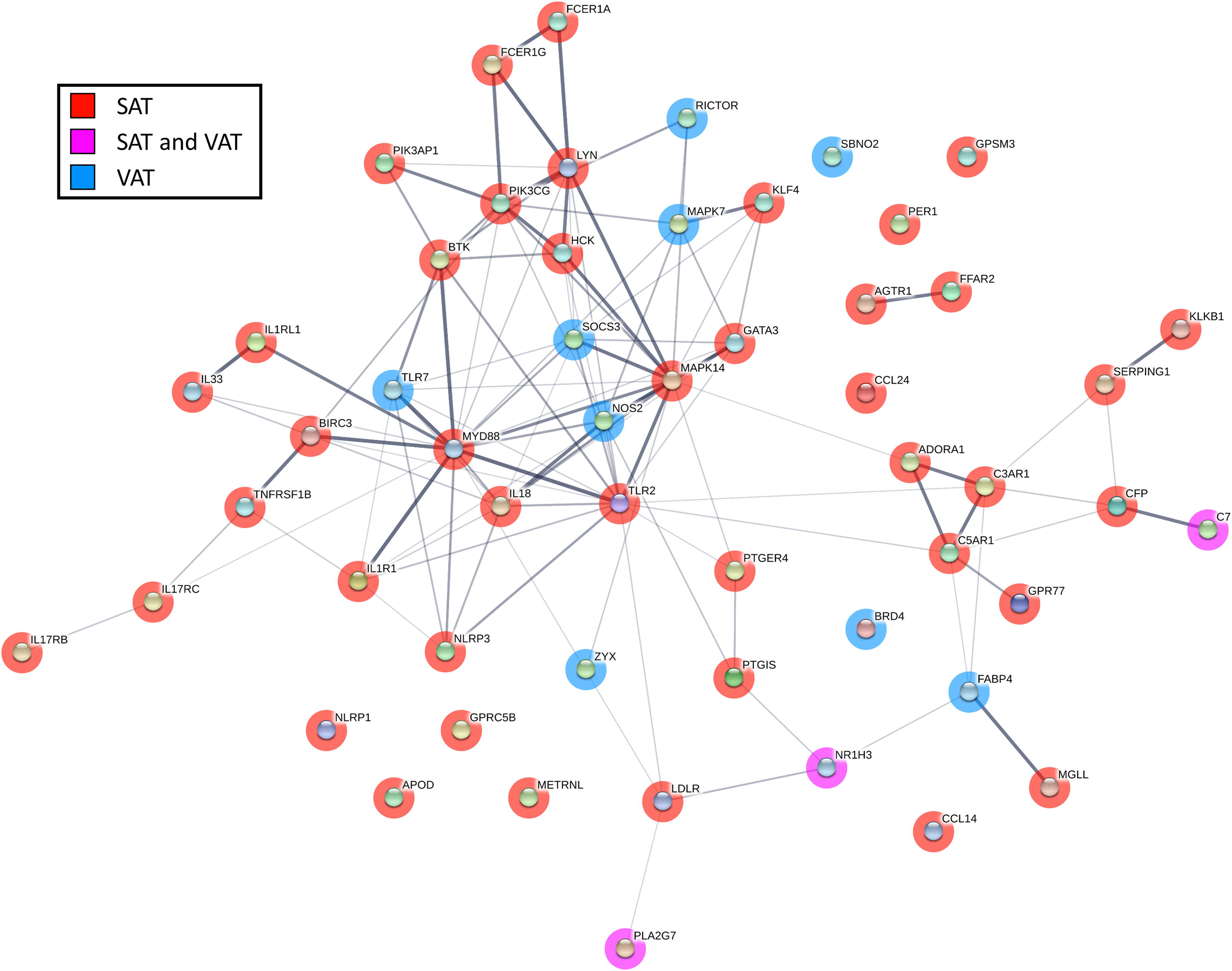
Network diagram of gene interactions in regulation of inflammatory response pathway, which was downregulated with prenatal bisphenol A treatment in subcutaneous adipose tissue (SAT) from 21-month old female sheep. Minimum interaction score of 0.3. Genes with p<0.05 shown as nodes. Genes in red have p<0.05 in SAT, blue have p<0.05 in visceral adipose tissue (VAT), and pink in both. Thickness of connection between nodes represent the level of evidence for interactions.

**Figure 4.**
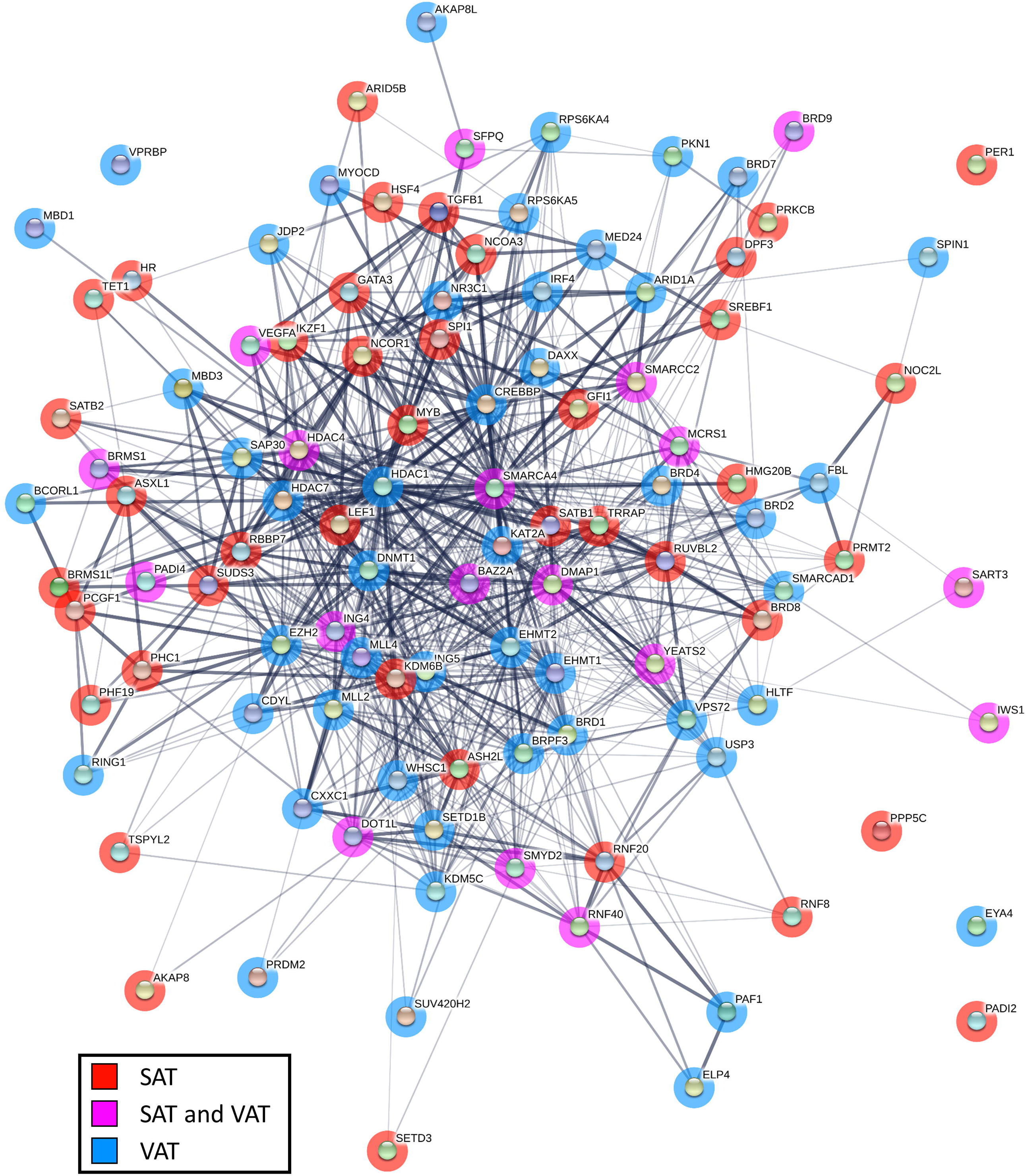
Network diagram of gene interactions in regulation of chromatin modification pathway, which was downregulated with prenatal bisphenol A treatment in visceral adipose tissue (VAT) and subcutaneous adipose tissue (SAT) from 21-month old female sheep. Minimum interaction score of 0.3. Genes with p<0.05 shown as nodes. Genes in red have p<0.05 in SAT, blue has p<0.05 in VAT, and pink in both. Thickness of connection between nodes represent the level of evidence for interactions.

## Discussion

The findings from this study demonstrate that SAT and VAT depots have distinct transcriptional profiles in sheep, similar to that observed in humans. Importantly, the results also show that prenatal BPA treatment induces depot-specific alteration of RNA expression of genes involved in inflammation, RNA processing, gene transcription, among others. These changes reflect the diverse roles of SAT and VAT in regulating lipid storage and insulin sensitivity, and indicate that adipose tissue transcriptional dysregulations may contribute to the metabolic disorders observed in prenatal BPA-treated female sheep. The implications of these findings are discussed below.

### Depot-Specific Transcriptional Changes in SAT and VAT

This study benchmarks physiologic gene expression differences between SAT and VAT in sheep. The divergent transcriptional profiles of SAT and VAT revealed by RNAseq analysis in control sheep is consistent with earlier reports in other species (Gesta et al., 2006; Hocking et al., 2010; Saely et al., 2012) and likely reflect the differences in their role in lipid storage, inflammatory status and distribution of thermogenic brown/beige adipocytes. As SAT favors lipid storage, it is not surprising that there was an increase in expression of genes such as HOXC9, HOXA9, IRX2 and engrailed homeobox 1 (EN1) that promote development and differentiation of white adipocytes (Yamamoto et al., 2010; Karpe and Pinnick, 2015), the adipose cell type specializing in lipid/triglyceride storage. These findings are also consistent with those reported in mouse SAT depots (Gesta et al., 2006; Yamamoto et al., 2010). The finding that SAT depots relative to VAT depots show increased expression of the transcription factor, T-box transcription factor 15 (TBX15), which participates in the development of thermogenic brown/beige adipocytes (Gburcik et al., 2012), is consistent with the observation that sheep SAT depots have higher expression of the thermogenic adipocyte marker, uncoupled protein 1 (UCP1) (Puttabyatappa and Padmanabhan, *unpublished*). These depot-specific gene expression profiles are in line with the SAT’s role in promoting metabolic homeostasis through sequestering lipids from entering circulation and deposition in ectopic locations (Frayn, 2002; Manolopoulos et al., 2010) and/or utilization of lipids by thermogenic adipocytes (Ishibashi and Seale, 2010).

Although VAT are also rich in white adipocytes, unlike SAT, VAT depots are less responsive to anabolic effects of insulin, they secrete proinflammatory cytokines and attract inflammatory cells (Hajer et al., 2008). These features make VAT lipolytically active, which promotes increases in circulating lipids and ectopic lipid deposition in insulin target tissues (Wajchenberg, 2000; Hajer et al., 2008; Bjorndal et al., 2011). Therefore, increased adiposity of VAT depots in particular is a risk factor for the development of metabolic disorders (Wajchenberg, 2000; Bjorndal et al., 2011). The findings that VAT was characterized by increased expression of the proinflammatory gene EPHX2 and adiposity associated transcription factor CUX1 are consistent with VAT physiology. Increased expression of EPHX2 has also been reported as a feature of porcine VAT (Gondret et al., 2012). Increases in EPHX2 expression would likely provoke an inflammatory state in VAT as it converts anti-inflammatory arachidonic acid metabolites into inactive diols (Newman et al., 2005). In other studies, silencing of this gene ameliorates high fat diet induced obesity associated complications in mouse (Bettaieb et al., 2013). The transcription factor CUX1 is involved in expression of FTO alpha-ketoglutarate dependent dioxygenase and retinitis pigmentosa GTPase regulator-interacting protein-1 like (RPGRIP1L) (Stratigopoulos et al., 2011), which is strongly associated with adiposity (Frayling et al., 2007; Baker and Beales, 2009). Although these gene expression patterns are consistent with the negative role VAT plays in regulating lipid storage and metabolic homeostasis, paradoxically genes involved in maintaining insulin sensitivity, specifically TF (McClain et al., 2018) and adipogenesis, such as glycerol-3-phosphate acyltransferase 3 (GPAT3) (Shan et al., 2010), were also upregulated in VAT in this study. As GPAT3 knockout mouse are protected from diet induced obesity (Cao et al., 2014), the increased expression of GPAT3 and CUX1 raises the possibility that VAT manifest a gene expression profile that favors increase in visceral adiposity under conditions of positive energy such as overfed state. In contrast, the increase in TF may be indicative of compensatory process in place to overcome the potential proinflammatory state to maintain insulin sensitivity of this depot, a possibility that needs to be tested further.

Comparing patterns in gene expression between SAT and VAT, gene set enrichment indicated genes involved in skeletal system morphogenesis were upregulated in SAT. While the reason for this increase is not clear, genes involved in skeletal system development are reportedly downregulated in VAT from diet-induced obese mouse (Choi et al., 2015). As this mouse model has increased adiposity with reduced insulin sensitivity and dyslipidemia (Luo et al., 2016), genes involved in skeletal system development may have a role in lowering risk of insulin resistance, consistent with the metabolic role of SAT relative to VAT (McLaughlin et al., 2011). In contrast, VAT showed increased enrichment of genes involved in toxic/xenobiotic metabolism, mitochondrial membrane and vasodilation. The increase in these genes may be a result of crosstalk between the gut, liver and VAT due to close proximity (Konrad and Wueest, 2014) and may stem from the fact that both gut and liver form the first line of defenses against xenobiotic and toxic substances (Hanninen et al., 1979). Likewise, upregulation of genes involved in vasodilation in VAT are consistent with the observation that visceral adiposity is associated with hypertension (Hall et al., 2015). While further roles of these depot-specific pattern of gene expression needs to be explored, these differences likely contribute to their differing functions in maintaining metabolic homeostasis.

### Effect of Prenatal BPA-Treatment on SAT Transcriptome

Prenatal BPA treatment induced marked changes in gene expression in the SAT depot of the adult sheep. Specifically, BPA treatment increased expression of multiple uncharacterized genes, ribosomal RNA, cyclin N-terminal domain containing 2 (CNTD2), transfer RNA lysine (anticodon CUU) (TRNAK-CUU_8), serglycin (SRGN), and serine/threonine kinase 17a (STK17A). Their roles in modulating SAT function remain to be determined. Gene network analysis also showed large number of genes modulated by prenatal BPA in SAT than VAT were related inflammation and immune functions. BPA treatment down regulated multiple gene sets related to immune function: leukocyte activation, migration, aggregation and cell-cell adhesion, lymphocyte activation and aggregation, T cell activation and aggregation, inflammatory response, cytokine production, positive regulation and activation of immune response (**Table 6**). Similarly, the physiologic basis for downregulation of genes involved immune function with BPA treatment in SAT is not known. Since the SAT depot maintains a low inflammatory profile and promotes healthy metabolic state (McLaughlin et al., 2011), these findings may reflect a compensatory processes in SAT to overcome systemic insulin resistance observed in prenatal BPA-treated sheep (Veiga-Lopez et al., 2016). Similar, albeit lack of changes in proinflammatory gene expression in SAT is also observed in sheep treated during the same susceptibility window with native steroid, testosterone, which also manifest peripheral insulin resistance (Puttabyatappa et al., 2017). Interestingly, prenatal testosterone-treated sheep have SAT that maintains responsiveness to insulin (Lu et al., 2016), suggesting of such a possibility in SAT from prenatal BPA-treated sheep.

### Effect of Prenatal BPA-Treatment on VAT Transcriptome

Prenatal BPA-treatment induced modest gene expression changes in the adult VAT depot. Specifically, BPA-treatment increased expression of transcriptional factors that participate in adipogenesis, such as AT-rich interaction domain 3A (ARID3A), zinc finger protein 565 (ZNF565) and RecQ like helicase 4 (RECQL4). These findings are consistent with the reports of increased adipogenesis by gestational BPA treatment in fetal sheep (Pu et al., 2017) and in BPA-treated human adipose stromal/stem cells (ASC), human omental fat cells, and 3T3-L1 preadipocytes (Sargis et al., 2010; Wang et al., 2013; Ohlstein et al., 2014). The zinc finger protein family is well known for its role in the promotion of adipogenesis and determination of adipogenic lineage (Wei et al., 2013), although the specific role of ZFP565 still needs to be elucidated. Both RECQL4 and ARID3A are involved in regulation of adipogenesis. They seem to however have contrasting roles as knockdown of RECQL4 reduces adipogenesis (Chacko, 2017), while depletion of ARID3A expression in ASC promotes adipocyte formation (Webb, 2008). As such, the net effect of BPA treatment on programming adipogenesis likely depends on the sum of effects of multiple transcription factors involved in BPA-induced adipogenesis in human preadipocytes (Boucher et al., 2014b).

Interestingly prenatal BPA-treatment induced upregulation of multiple small nucleolar RNA levels (LOC114113403, LOC114117716 and LOC114116238). These are noncoding RNAs, transcribed by RNA Polymerase II and are usually involved in processing and modification of other RNAs and can occasionally act like microRNA to induce epigenetic alterations (Ender et al., 2008; Kufel and Grzechnik, 2019). Consistent with the increase in these small nucleolar RNA transcripts, gene network and gene enrichment analysis revealed BPA treatment upregulated multiple gene sets related to RNA splicing, such as mRNA processing, negative regulation of transcription from RNA polymerase II promoter, transcription factor activity, binding and cofactor activity, and RNA splicing. In addition, BPA treatment upregulated gene sets involved in chromatin, histone and peptidyl-lysine modification that were enriched specifically in VAT than SAT depot. These changes are likely indicative of a role for prenatal BPA treatment induced epigenetic modification in these depots. These results are consistent with reported associations between epigenetic changes involving both histone modification and noncoding RNA expression with BPA exposure as a mechanism through which BPA programs its effects (Alonso-Magdalena et al., 2016).

### Conclusions

The findings from the present study on impact of prenatal BPA exposure on depot-specific transcriptional profile is of clinical and public health significance in view of the fact that pregnant women are ubiquitously exposed to BPA (Woodruff et al., 2011) and BPA has a role in promoting adipogenesis and obesity (Thayer et al., 2012; Lubrano et al., 2013). Evidence to date point to EDC contribution to the significant increases in incidences of adipose defects associated with conditions such as abdominal obesity (Trasande et al., 2012; Liu et al., 2017). The finding that prenatal BPA exposure highly upregulates genes associated with adipogenesis and adiposity in VAT raises the possibility that BPA’s contribution to promoting obesity may be VAT depot-specific. While it is well accepted that epigenetic reprogramming forms the basis of BPA-associated adult onset diseases, for the most part such evidence in human tissues comes from non-adipose tissues (Doherty et al., 2010). Evidence from the present study pointing to predominant changes in genes involved in chromatin modification in both VAT and SAT, is in line with epigenetic contributions to adipose defects. However, these gene sets are highly upregulated in the VAT (**Supplemental Table 7**), a depot with higher risk for development of obesity. Importantly, the pathways impacted by prenatal BPA treatment, such as the inflammatory and epigenetic pathways, provide targets for developing interventions to prevent prenatal BPA-induced pathology from emerging.

A major strength of this study is the assessment of the depot-specific transcriptional changes using the robust RNAseq approach in an animal model of translational significance. The human translational significance of the sheep model relates to species similarity in fetal developmental timeline, precociality at birth, the homology in adipogenesis and distribution of brown/beige fat (Padmanabhan and Veiga-Lopez, 2014; Gonzalez-Bulnes and Chavatte-Palmer, 2017; Fuller-Jackson and Henry, 2018), and development of insulin resistance with adiposity (Clarke, 2008). Importantly these data were generated utilizing a BPA dose that resulted in fetal exposure typical of environmental exposure to a pregnant woman (Gerona et al., 2013; Veiga-Lopez et al., 2013a; Veiga-Lopez et al., 2015b; Lee et al., 2018). These findings from this study should also be interpreted taking into account the limitation - the assessment of transcriptome profile in whole adipose tissues; adipose tissues are made up of diverse cell types including adipocytes, immune and vascular cells. Nonetheless, the findings are in agreement with the dimorphic role of VAT and SAT on lipid metabolism and insulin sensitivity lending support on the validity of the experimental approach. Other limitations include lack of inclusion of interventional studies to overcome the pathology, an aspect to be considered in future investigations. In conclusion, the results from this study show that exposure to prenatal BPA has adipose-depot specific impact on genes involved in inflammation, adipogenesis, adipocyte differentiation, and chromatin remodeling and provide mechanistic cues that could be exploited to understand the pathogenesis and in developing therapeutic strategies.

## Acknowledgements

We thank Mr. Douglas Doop and Gary McCalla for their valuable assistance in breeding, lambing, and careful animal care; Dr. Almudena Veiga-Lopez, Dr. Bachir Abi Salloum, Mr. Evan Beckett, Mrs. Carol Herkimer and students supported through the Undergraduate Research Opportunity Program (University of Michigan) for the help provided with administration of treatments and tissue harvest.

Research reported in this publication was supported by R01-ES-016541 (VP) and P30 ES017885. KB and JD are supported by R01 ES025574, R01 ES025531, R01 MD013299, R01 AG055406. MP is supported via Ruth L. Kirschstein Institutional Training Grant T32 ES007062.

## Disclosure statement

Authors have nothing to disclose.

