## Supplementary Material for "Developmental programming: Transcriptional regulation of visceral and subcutaneous adipose by prenatal bisphenol-A in female sheep"

**Supplemental Table 1**. Sample sequencing descriptive statistics. Number of reads per sample differed by tissue type (t-test p-value = 8.1x10^-5^), but not by BPA treatment (t-test p-value = 0.35). Number of genes per sample did not differ by tissue type (t-test p-value = 0.39), but did differ by BPA treatment (t-test p-value = 0.01).

| **Sample** | **Adipose Deposit** | **Treatment** | **N Reads** | **N Genes** |
| --- | --- | --- | --- | --- |
| 124151 | Visceral | Control | 11639272 | 18914 |
| 124152 | Visceral | Control | 10998858 | 18480 |
| 124153 | Visceral | Control | 12037279 | 18665 |
| 124154 | Visceral | Control | 13110261 | 18007 |
| 124155 | Visceral | BPA | 9604422 | 18084 |
| 124156 | Visceral | BPA | 9035630 | 18022 |
| 124157 | Visceral | BPA | 6768945 | 16926 |
| 124158 | Visceral | BPA | 14305764 | 17951 |
| 124159 | Subcutaneous | Control | 6223460 | 18481 |
| 124160 | Subcutaneous | Control | 5269312 | 18390 |
| 124161 | Subcutaneous | Control | 6193370 | 18122 |
| 124162 | Subcutaneous | Control | 4738482 | 17509 |
| 124163 | Subcutaneous | BPA | 4828287 | 17751 |
| 124164 | Subcutaneous | BPA | 4314984 | 17619 |
| 124165 | Subcutaneous | BPA | 3046126 | 17487 |
| 124166 | Subcutaneous | BPA | 4349021 | 17884 |

**Supplemental Table 2**. Results for differential gene expression by adipose tissue type, comparing subcutaneous control to visceral tissue control.

File: DE SAT vs VAT.csv

**Supplemental Table 3**. Gene set enrichment results from RNA-enrich for adipose type differences comparing subcutaneous to visceral fat

File: RNA-Enrich_tissue.csv

**Supplemental Table 4**. Results for differential gene expression by BPA exposure, comparing subcutaneous exposed to subcutaneous control group.

File: BPA (subcutaneous) DE.csv

**Supplemental Table 5**. Results for differential gene expression by BPA exposure, comparing visceral exposed and control groups.

File: BPA (visceral) DE.csv

**Supplemental Table 6**. Gene set enrichment results from RNA-enrich for BPA in subcutaneous fat

File: RNA-Enrich_bpa_sat.csv

**Supplemental Table 7**. Gene set enrichment results from RNA-enrich for BPA in visceral fat

File: RNA-Enrich_bpa_vat.csv

**Supplemental Figure 1**. Plot of first two principal components calculated on variance stabilizing transformed count data. Points of plots are colored by treatment group and adipose tissue type. Groups, each with four samples, were BPA treated subcutaneous fat, control subcutaneous fat, BPA treated visceral fat, control visceral fat.

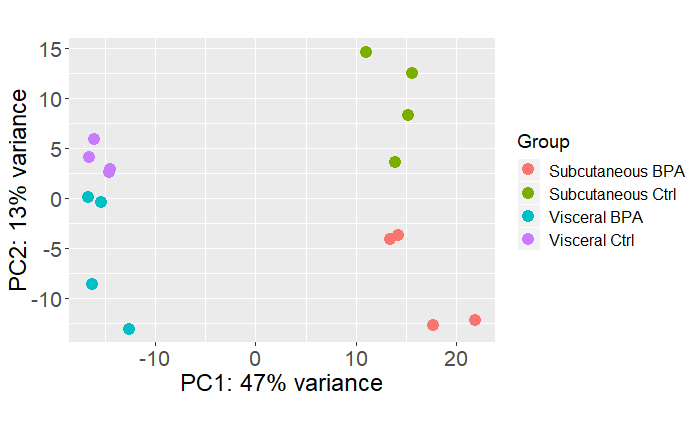

**Supplemental Figure 2A**. Normalized counts, plotted on log_2_ scaled coordinates, of *HOXC9* gene in control samples. Y-axis is log_2_ scaled. **Figure 2B.** Normalized counts, plotted on log_2_ scaled coordinates, of *IRX2* gene in control samples.

| **A.** 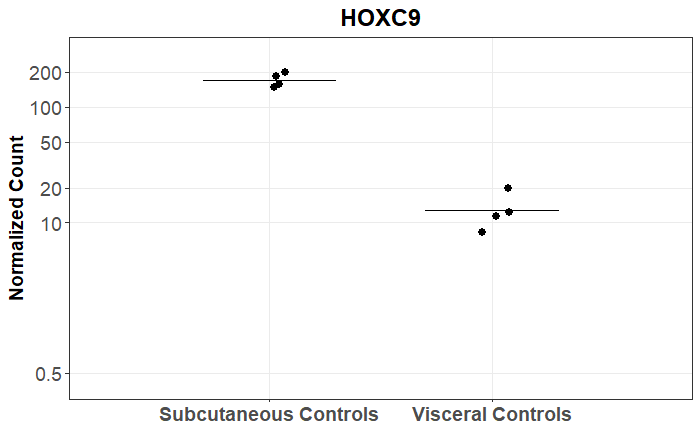 | **B.**  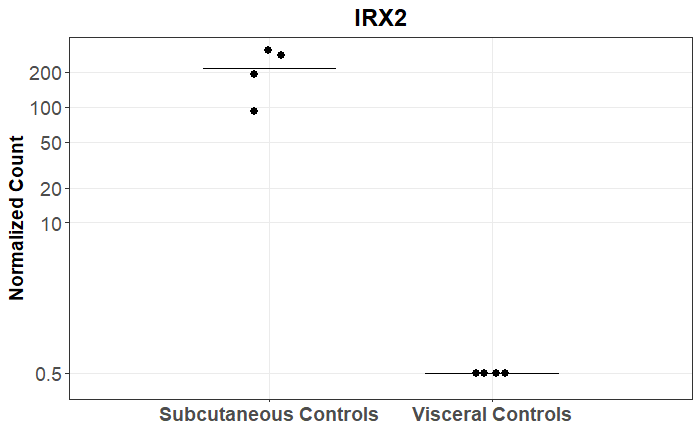 |
| --- | --- |

**Supplemental Figure 3A**. Normalized counts, plotted on log_2_ scaled coordinates, of *CNTD2* gene in subcutaneous fat samples. **Figure 3B.** Normalized counts, plotted on log_2_ scaled coordinates, of *LOC114112289* gene in subcutaneous fat samples.

| **A.**  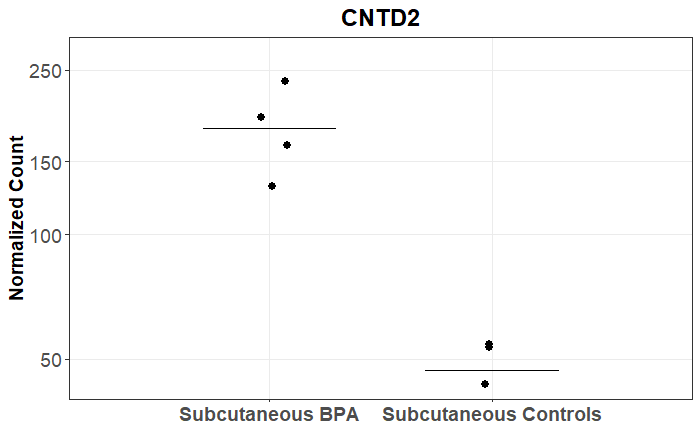 | **B.**  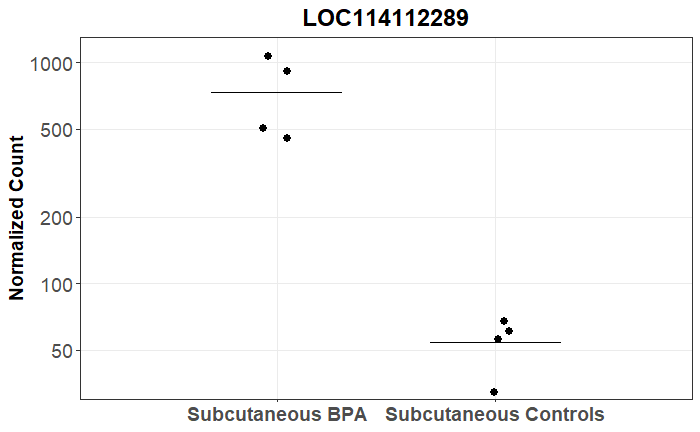 |
| --- | --- |

**Supplemental Figure 4.** Heatmaps of individual sample gene expression, separated by cell type. For each heatmap, the top 200 genes differentially expressed with BPA were used.
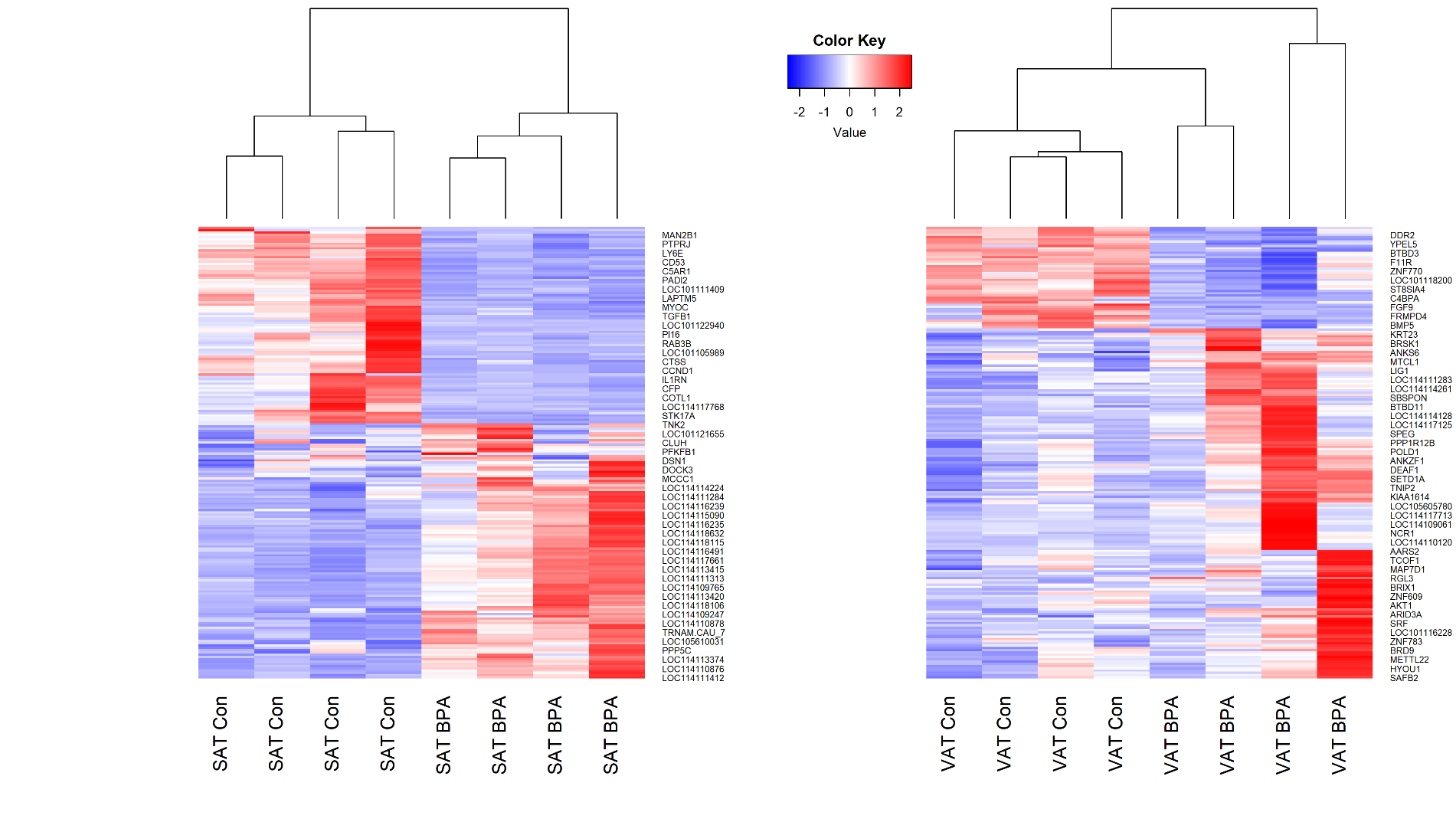
